# Cryoprotectants-assisted plunge freezing of thick brain tissue specimens for targeted physiologically relevant cryo-imaging *in situ*

**DOI:** 10.1101/2025.10.09.681493

**Authors:** Ash Weier, Louis Perez, Fei Gao, Elliot T. Morgan, Pingting Liu, Ivan C. Mounteer, Garry P. Morgan, Qian Shi, Fabio A. Vigil, Lydia-Marie Joubert, Andreas Hoenger, Michael H. B. Stowell, Oleg Klykov

## Abstract

*In situ* cryoET (cryoelectron tomography) and cryo-FIB/SEM (cryo-focused ion beam/scanning electron microscopy) volume-EM (electron microscopy) imaging provide spatiotemporal snapshots of biological systems in their near-native aqueous environment. Freezing and subsequent thinning of thick biological specimens prior to cryo-imaging is a time-consuming and challenging task that requires state-of-art methodology. As a result, cryo-imaging reports obtained from non-trivial specimens including mammalian brain tissues are scarce and their physiological relevance remains to be determined. Here, we benchmarked plunge freezing with a variety of cryoprotectants that allow for mouse brain tissue vitrification of up to about 100 microns thick and across several brain regions while keeping the tissue functional. By utilizing the knock-in (KI) mouse model with fluorescent astrocytes we have performed targeted cryo-FIB/SEM volume-EM imaging as well as targeted high-resolution cryoET imaging. Prior to cryoET, we have successfully generated lamellae in a semi-automated fashion on both LMIS (liquid metal ion source)- and plasma-based cryo-FIB/SEM instrumentation thus expanding applicability of our pipeline. We visualized the NVU (neurovascular unit) and astrocyte’s processes and validated the physiological relevance of our outputs based on the morphology of the corresponding cellular and subcellular features. The pipeline utilizes common vitrification setups and can be potentially extended toward alternative tissue specimens. Ultimately, we expect our approach to become an important step towards democratization of physiologically relevant *in situ* cryo-imaging studies.

## Introduction

Understanding biological processes in physiologically relevant aqueous environments is vital for obtaining actionable translational insights. Specifically, accessing protein structure and interactions coupled with cellular and organelle morphology within brain tissue under physiological conditions is critical for unraveling the mechanisms underlying neurological functions and for developing novel therapeutic targets against neurological disorders. Large-volume cryo-FIB/SEM tomoapplication (cryo-Focused Ion Beam/Scanning Electron Microscopy, cryo-volume-EM, or cryo-Serial Surface Imaging) can be utilized to access the contextual information including cellular morphology and interactions between the cells or organelles (Collinson et al., 2023). In contrast, cryoET (cryoelectron tomography) has emerged as a powerful tool for visualizing proteins and protein complexes at molecular and residue-level resolution and with minimal interference of biological function (Beck et al., 2007; Grünewald et al., 2003; Mahamid et al., 2016; Tegunov et al., 2021). Unlike traditional methods that typically isolate proteins and protein ensembles from their physiological context, cryoET, combined with cryo-focused ion beam (FIB)-milling and subtomogram averaging (STA), enables high-resolution three-dimensional visualization of protein machinery in their near-native environments (Marko et al., 2007; Noble and De Marco, 2024).

Cryo-imaging pipelines on tissues can be performed by specimen sectioning with commonly available vibratomes. A thickness of 20 microns is readily achievable for specimens that have been pre-fixed for immunohistochemical staining, while obtaining slices thinner than 100 microns from fresh tissue is challenging and requires an experienced operator. Vibratome sectioning of fresh tissue frequently results in surface layer damage due to shearing forces, which necessitates extended recovery times, and leaves only a narrow band of physiologically relevant fresh tissue suitable for cryo-imaging (Suter et al., 1999). While agarose embedding facilitates fresh tissue thinning (Creekmore et al., 2024), it may cause compression of specimens and surface layer damage, potentially compromising physiologically relevant outputs. Additionally, the dense shell can also hinder oxygen and substrate diffusion to inner cell layers, further affecting tissue viability and function during physiological assessment (Bussek et al., 2012). Concurrently, specimens must maintain sufficient thickness to retain a sufficient number of regions of interest (ROIs) representing relatively rare molecular or physiological events. Thus, optimal tissue thickness lies between 50-100 microns, which balances structural integrity and the ability to identify target features.

Next, cryo-preservation of samples through vitrification is performed and locks protein machinery in its near-native, *in situ*, hydrated state. While the vitrification of cellular samples is relatively straightforward and can be done with conventional plunge freezing (Mahamid et al., 2016; Medalia et al., 2002; Chaikeeratisak et al., 2019; Allegretti et al., 2020), thicker samples such as tissues are typically vitrified with High Pressure Freezing (HPF) (Harapin et al., 2015). The Waffle method (Kelley et al., 2022) offers an efficient and high-throughput HPF-based approach that eliminates preferred orientation, is performed on-grid, and allows for using high sample concentrations. For vibratome-produced tissue specimens, the underlying geometry of the grid bars does not typically allow for the specimen to be frozen with conventional Waffle approach, however the modified Waffle method has been successfully implemented on mouse hippocampal tissue slices (Matsui et al., 2024). For previously-frozen human brain autopsies, HPF has already provided potentially translationally-actionable insights, but interference of crystalline ice with the native environment requires further investigations (Gilbert et al., 2024). Conventional HPF typically involves specialized carriers and additional fillers like 2-methylpentane that increase sample thickness, tissue thinning time for subsequent cryoET imaging, and complicates detection of the ROIs when no fluorescent label is present.

A viable vitrification alternative, plunge freezing (PF), has emerged as a standard vitrification method for samples less than 10 microns thick. PF requires rapid heat transfer which is essential for achieving a successful vitreous state (Adrian et al., 1984; Dubochet et al., 1988). Thicker samples pose a challenge, as slower heat transfer may hinder vitrification and lead to ice crystal formation within the sample. Recently, several reports on cryoprotectant-assisted on-grid PF of thick specimens have demonstrated its feasibility for vitrifying brain tissues and thicker cells. Specifically, *Drosophila* brains were successfully vitrified after an incubation in 10% glycerol (Bäuerlein et al., 2021). For mammalian oocytes, vitrification was achieved via a gradient of dimethyl sulfoxide (DMSO) and ethylene glycol (EG) (Jentoft et al., 2023). For human brain tissue, reported on-grid vitrification requires prolonged incubation in a solution containing 20% glycerol and 1 M trehalose coupled with agarose-embedding before tissue slicing (Creekmore et al., 2024). None of the reports above have examined the physiological relevance of their outputs, nor the effect of cryoprotectants on the sample. In the case of trehalose, for example, the water replacement hypothesis postulates that trehalose’s hydroxyl groups substitute for water in an aqueous protein environment (Stolz et al., 2011), distortions and making structural insights less relevant. Most importantly, while no evidence of osmotic effects was observed, the usage of cryoprotectants has the potential to cause cellular damage when the osmolarity is significantly higher compared to physiological conditions (Ballyk et al., 1991a; Elliott and Jasper, 1949). Thus each tissue type requires custom, sample-tailored vitrification protocols that involve meticulous optimization of chemicals utilized for successful and physiologically relevant PF of thick specimens.

For cryo-FIB/SEM volume-EM imaging, vitrified samples of any thickness can be imaged, but for high-resolution cryoET and STA, thin sample slices with a thickness below 200-300 nanometers are required (Neselu et al., 2023). Cryo-FIB sample thinning is currently the method of choice for creating electron-transparent sample slices (lamellae) from PF cellular samples that are below 10 microns thick (Marko et al., 2007; Schaffer et al., 2015). For thicker HPF samples, sophisticated, time-consuming, and ice-contamination prone strategies based on conventional cryo-lift out (cryo-LO) are preferred, but they require additional hardware, custom-made cryo-EM grids, and significant operator expertise, thus making cryo-LO methods limited and only available to highly specialized laboratories (Klumpe and Erdmann, 2024; Rubino et al., 2012; Schaffer et al., 2019). At the same time, on-grid lamellae preparation is done with the basic cryo-FIB/SEM instrumentation and has been proven successful even for thick specimens (Bäuerlein et al., 2021; Creekmore et al., 2024; Harapin et al., 2015). The main disadvantage of on-grid thinning is a loss of the vertical sample dimension and this information is not preserved for high-resolution cryoET imaging as opposed to modern serial versions of cryo-LO (Nguyen et al., 2024; Schiøtz et al., 2023). Otherwise, on-grid lamellae preparation is straightforward and can be performed in a semi-automated fashion similar to advanced cryo-LO implementations like e.g. SOLIST (Nguyen et al., 2024).

Here we developed a chemically assisted and physiologically relevant PF-based solution for cryo-imaging of thick mouse brain specimens that is readily adoptable by the majority of cryo-EM laboratories. Because most disease models are based on transgenic mice, we have benchmarked PF for thick samples using mouse hippocampal, cortical, and subregions of striatum brain tissue slices generated with a common vibratome setup. We then tested several cryoprotectant mixtures that ensure sufficient tissue vitrification and physiological relevance of cryo-imaging outputs. To assess vitrification, we prepared a set of lamellae placed at different distances from the surface of brain slices of various thicknesses and then estimated the quality of vitrification within those lamellae via cryoET. The physiological relevance was evaluated by comparison of morphological features visualized by a targeted cryo-FIB/SEM volume-EM and cryoET imaging against published room temperature (r.t.) EM data. Specifically, by utilizing a knock-in (KI) mouse model with astrocytes expressing tdTomato, we imaged components of a neurovascular unit (NVU) with cryo-FIB/SEM volume-EM imaging and demonstrated expansion (Glynn et al., 2024) of the corresponding cellular features in comparison to r.t. EM. With targeted cryoET, we visualized subcellular features specific for astrocyte processes and validated their physiological relevance. We have also benchmarked semi-automated on-grid sample thinning of thick specimens on both LMIS (Liquid Metal Ion Source)- and plasma-based (pFIB) instrumentation. While every organ within an organism may require an optimization of the cryoprotectant mixture to ensure vitrification and physiological relevance of PF outputs, the lamella preparation strategy is universal and potentially can be extended towards organs other than the brain. By democratizing cryo-imaging aimed at determination of physiologically relevant cellular morphology and structure of the proteins in their near-native context, our findings help to bridge the gap between structural biology and translational investigations of brain function, thus advancing research in neurological science.

## Results

The main goal of this study is to determine the most efficient cryoprotectants mixture that allows for reliable vitrification with PF while yielding cryo-imaging outputs usable for translational research. This task develops a robust, easy-to-implement, widely accessible pipeline for subsequent cryo-EM imaging. In general, HPF-based approaches are considered more robust for thicker specimens in terms of vitrification as opposed to PF for which a conventionally accepted limit is 10 microns. At the same time, HPF implies utilization of specialized carriers that complicate subsequent sample thinning. Additionally, navigation through the sample prepared with a spacer or fillers like 2-methylpentane, dextran or BSA (bovine-serum-albumin) is not straightforward especially without previous ROI definition by cryo-FLM (cryo-Fluorescent Light Microscopy). HPF instrumentation is also not as available as PF setups that are utilized for single particle analysis (SPA) pipelines and are common in cryo-EM facilities.

Successful PF of thick brain specimens has been reported before, however, there is no information available regarding the physiological relevance of the corresponding outputs. Thus, our first task was to find the most appropriate combination of chemicals that 1) do not interfere with the physiological state of the mouse brain tissue and 2) ensure the specimen’s suitability for cryo-transmission electron microscopy (cryo-TEM) with imageable quality of vitrified ice. The utilized aCSF solution had reduced Ca^2+^ levels thus minimizing cell death signaling. We have accessed several combinations of chemicals, both new and previously reported, and after developing a FIB-milling protocol that is easily reproducible, semi-automated, and works on both new and previous generation of FIB/SEM setups, we were able to find the most appropriate cryoprotectant mixture that robustly provides a medium to high quality vitrification.

We then utilized KI mice models with fluorescent astrocytes and imaged an NVU with cryo-FIB/SEM volume-EM as well as astrocyte’s processes with cryoET. Astrocyte processes exhibit characteristic tube-like motifs driven by astrocytes’ function to wrap around blood vessels (Steinman et al., 2017) making them an easily detectable target even within the thick tissue specimens. Targeted cryo-imaging allows for comparison of the morphology of the cellular features such as pericytes, endothelial layers, and astrocyte endfeet wrapping around a vascular unit to the previously reported r.t. EM data. Thus, obtained cryo-imaging outputs confirm that our pipeline is providing the physiologically relevant outputs. The generic workflow is described in the **Supplementary Figure 1**.

### Thick specimen vitrification with PF and on-grid thinning

Prior to PF, we have soaked the samples in artificial cerebrospinal fluid (aCSF) with a mixture of cryoprotectants. To maintain the physiologically relevant environment after adding cryoprotectants, we assessed how the osmolarity is affected in comparison to PBS or aCSF without additives (**Table 1**). In the cases of the most commonly used chemicals for PF such as glycerol with and without trehalose, the osmolarity is at least 4-fold (without trehalose) and 6-fold (with 0.3 M trehalose) higher than acceptable range of aCSF. The addition of polyvinylpyrrolidone (PVP) does not exhibit potentially debilitating effects of the glycerol and allows for maintaining the osmolarity within acceptable values. Specifically, 10% PVP with 0.3 M trehalose was measured as 608 mOsm/kg and 15% PVP as 352 mOsm/kg thus placing those solutions as 2-fold above the acceptable limit and approximately within the desired range respectively.

**Table 1.** Physiological relevance of the solutions with cryoprotectants based on the osmolarity values. Values represent the average osmolarity among technical triplicates of solutions with the specified compositions. With the typical aCSF osmolarity of ∼310 mOsm/kg (EllioÄ and Jasper, 1949; Oliveira et al., 2021; Segev et al., 2016), we define an acceptable range of 300-350 mOsm/kg (Ballyk et al., 1991b). Solutions highlighted in green represent a physiologically representative environment for mouse brain tissue slices and yellow indicate solutions no more than 2-fold of the acceptable physiological range.

|  |  |  |  |  |  |  |  |  |  |  |  |  |  |  |  |  |  |  |  |  |  |
| --- | --- | --- | --- | --- | --- | --- | --- | --- | --- | --- | --- | --- | --- | --- | --- | --- | --- | --- | --- | --- | --- |
| **Glycerol (v/v), %** | 20 | 20 | 15 | 15 | 15 | 10 | 10 | 10 | 0 | 0 | 0 | 0 | 0 | 0 | 0 | 0 | 0 | 0 | 0 | 0 | 0 |
| **PVP (w/w), %** | 0 | 0 | 0 | 0 | 0 | 0 | 0 | 10 | 10 | 10 | 0 | 0 | 0 | 0 | 0 | 20 | 20 | 15 | 15 | 10 | 10 |
| **Trehalose, M** | 1 | 1 | 0 | 0 | 0.3 | 0 | 0.3 | 0 | 0.3 | 0 | 0 | 0 | 0 | 0 | 0 | 0 | 0.3 | 0.3 | 0 | 0.9 | 0.6 |
| **Dextran, %** | 0 | 0 | 0 | 0 | 0 | 0 | 0 | 0 | 0 | 0 | 20 | 20 | 0 | 10 | 0 | 0 | 0 | 0 | 0 | 0 | 0 |
| **Sucrose, %** | 0 | 0 | 0 | 0 | 0 | 0 | 0 | 0 | 0 | 0 | 0 | 5 | 0 | 5 | 0 | 0 | 0 | 0 | 0 | 0 | 0 |
| **Ethylene Glycol, %** | 0 | 0 | 0 | 0 | 0 | 0 | 0 | 0 | 0 | 0 | 0 | 0 | 0 | 5 | 7.5 | 0 | 0 | 0 | 0 | 0 | 0 |
| **DMSO, %** | 0 | 0 | 0 | 0 | 0 | 0 | 0 | 0 | 0 | 0 | 0 | 0 | 0 | 0 | 7.5 | 0 | 0 | 0 | 0 | 0 | 0 |
| **Solvent** | aCSF | PBS | aCSF | PBS | PBS | aCSF | aCSF | aCSF | aCSF | aCSF | aCSF | aCSF | aCSF | aCSF | aCSF | aCSF | aCSF | aCSF | aCSF | aCSF | aCSF |
| **Osmolarity (mOsm/kg)** | 3404 | 2742 | 1440 | 843 | 3223 | 1824 | 2068 | 1866 | 644 | 318 | 365 | 603 | 294 | 1395 | 2750 | 405 | 823 | 756 | 352 | 1406 | 1077 |

To monitor the quality of vitrification, we have utilized an on-grid thinning approach followed by cryoET imaging (**Figure 1**). Direct on-grid milling has already been implemented to brain tissues and represents an attractive alternative to cryo-LO-based methods. At the same time, on-grid milling suffers from low throughput, lack of automation, and has not yet delivered a high-resolution, subnanometer structural output. This is largely explained by the methodological focus of the previous studies rather than benchmarking STA on trivial ribosome-like proteins. We have combined the front surface geometry (Creekmore et al., 2024) with deepFIB approach(Jentoft et al., 2023) and a semi-automated pipeline previously tailored for FIB-milling of Waffle specimens with both LMIS- (**Figure 1 H-K**) and plasma-based FIB/SEM instruments (**Figure 1 L-O**). The resulting pipeline takes approximately 359 minutes for LMIS and 76 minutes for pFIB setup per single lamellae of 120-200 nanometers thick as defined in AutoTEM Cryo. Manual site preparation takes the most of the time in case of LMIS and is reduced from 258 to about 11 minutes with pFIBs (with optimized current and gas species) while automated thinning takes only about 100 minutes per site for LMIS and 65 minutes for pFIBs.

**Figure 1.**
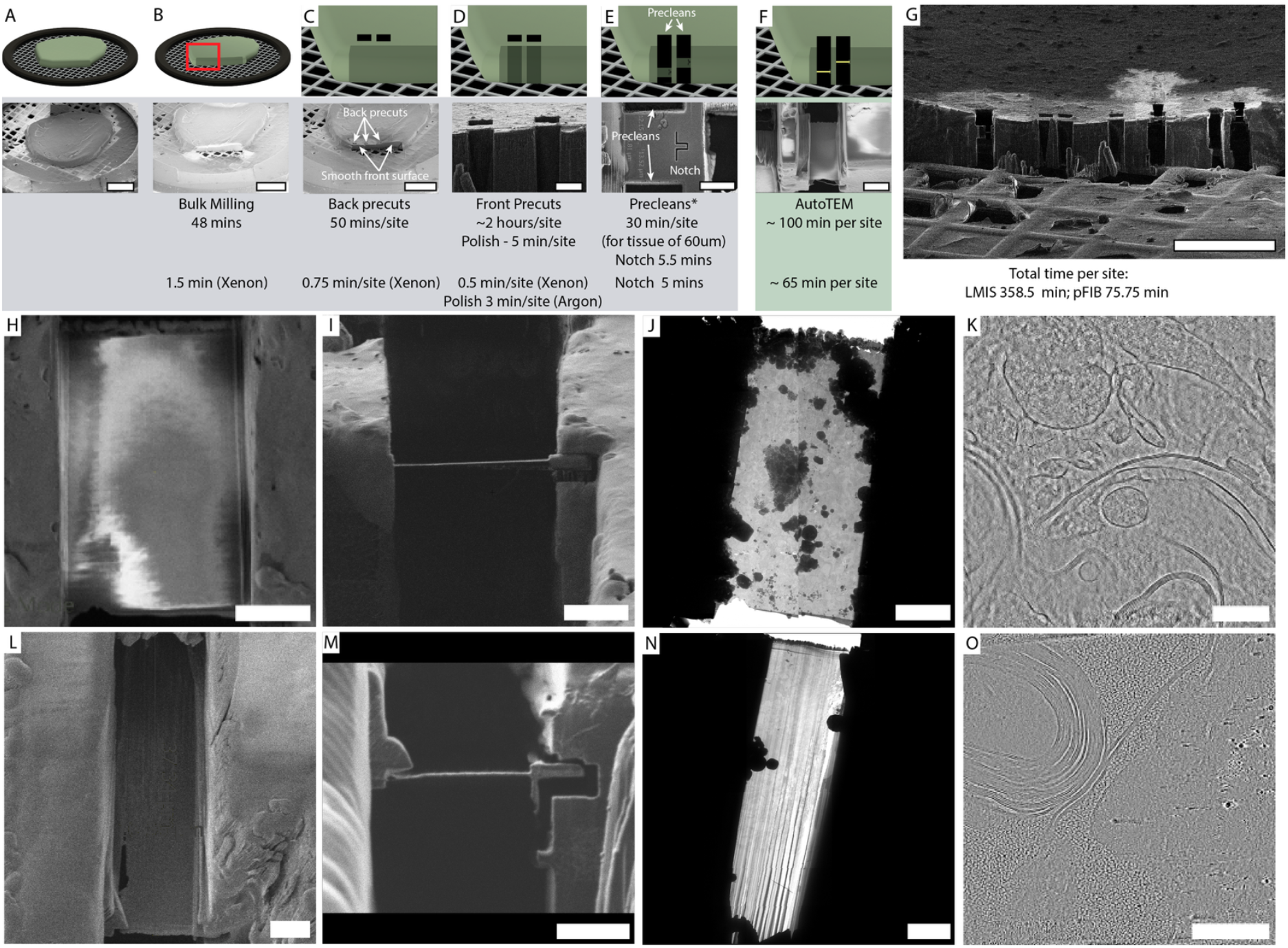
Lamellae generation for thick plunge frozen tissues. (**A**) SEM overview of 1.5 mm biopsy punch from mouse cortex before coating. (**B**) After Pt sputtering and GIS application, bulk milling is performed at higher currents to prepare the front surface. Typically, removal of 65 x 680 microns of tissue takes about 48 minutes with LMIS at 60 nA currents. (**C**) Back precuts and additional removal of 25 µm is performed at 15-16 nA. (**D**) Final front surface polishing is done at 1 nA and removes about 1 µm of the material. (**E**) Next, precleans and notch milling are performed on the front sample surface. Final slab thickness before automated lamellae preparation is 25-50 µm. (**F**) Final lamellae preparation with AutoTEM Cryo down to nominal thickness of 120-180 nm. (**G**) Overview of all lamella sites post milling. (**H-J**) and (**L-N**) show lamellae prepared with LMIS and pFIB respectively that were imaged in (**H, L**) SEM, (**I, M**) IB, (**J, N**) TEM. (**K**) and (**O**) show representative tomograms for LMIS and pFIB. Scale bars for (**A-C**) 400 µm, (**D)** 25 µm, (**E, F**) 10µm, (**G**) 150 µm, (**H-J**) 5 µm, (**K, O**) 200 nm, (**L-N**) 5 µm.

We have assessed ice quality across 15 lamellae from thick mice tissues soaked in 15% glycerol, 10% PVP, 10% PVP with 0.3 M trehalose, and 15% PVP (**Figure 2, Supplementary Figure 2, Supplementary Videos 1-6**). While 15% glycerol allows for generation of fully vitreous lamellae (**Figure 2A**), we were able to locate the artifactual “bent” features that are likely a consequence of the several-fold increased osmolarity of the solution (**Supplementary Figure 3**). 10% PVP with 0.3 M trehalose provides moderate but still imageable ice quality (**Figure 2B**) while 10% PVP alone results in multiple crystalline ice inclusions (**Figure 2C**). 15% PVP results in a fully vitreous lamellae while maintaining acceptable osmolarity values (**Figure 2D)**. At the same time, tomograms collected from samples vitrified with 15% PVP show a characteristic mesh throughout the tomogram and we interpret this mesh as an artifact cause by high PVP concentration.

**Figure 2.**
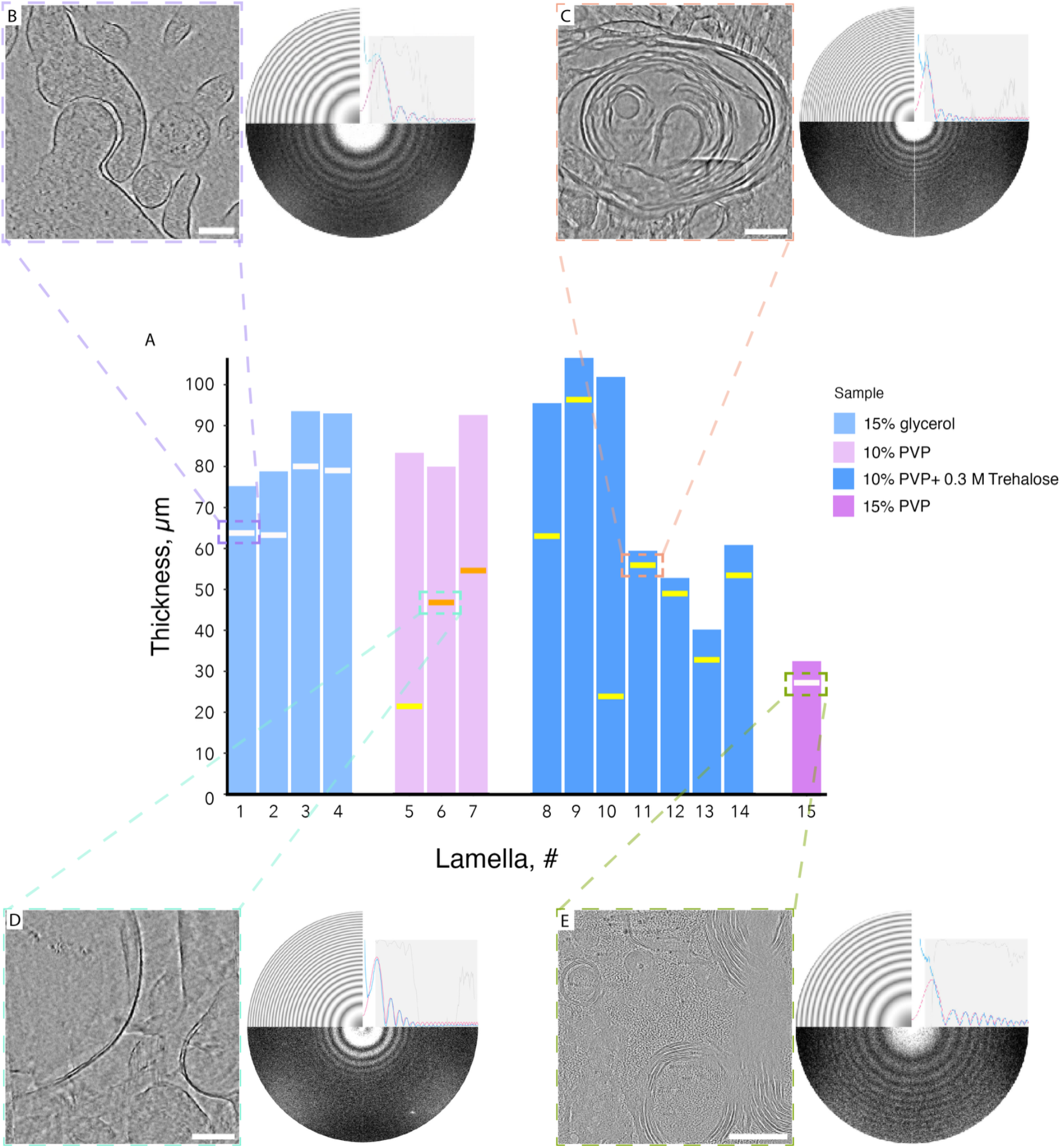
Plunge freezing efficiency across thick brain specimens. **(A)** Barplots representing local thickness of plunge frozen tissue slices. Lamellae position is shown by horizontal lines with color scheme based on the vitrification efficiency: white for fully vitreous, yellow for mostly vitreous with isolated crystalline ice inclusions present, and orange for partially vitreous with numerous crystalline ice inclusions. (**B-E**) - central slices through the tomograms for each condition with the corresponding FFTs. All samples are prepared in aCSF with addition of (**B**) 15% glycerol, (**C**) 10% PVP with 0.3 M trehalose,(**D**) 10% PVP, and (**E**) 15% PVP. All scale bars are 200 nm.

**Figure 3.**
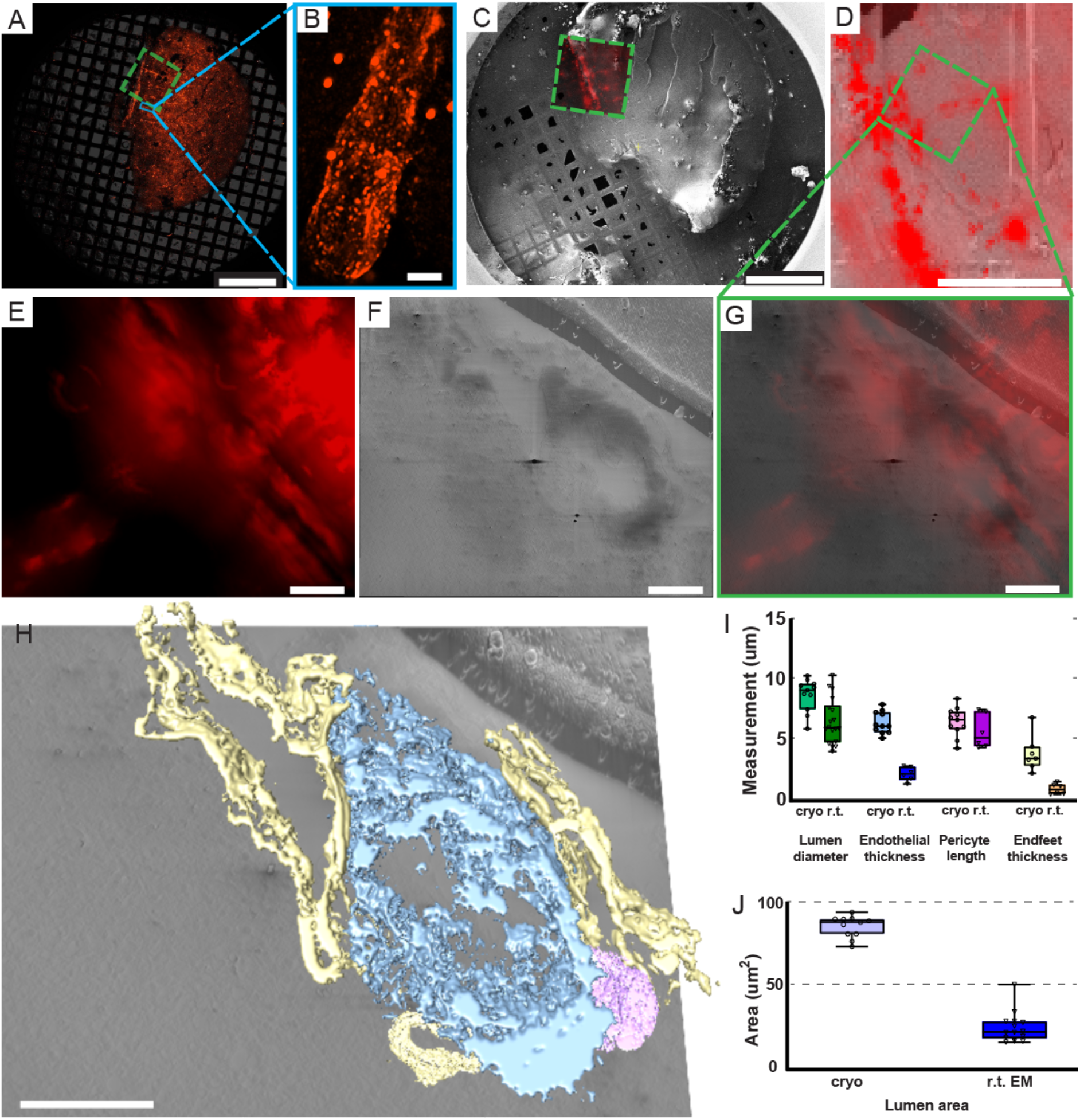
Fluorescence-guided targeting and cryo-FIB/SEM of mice NVU compartments with tdTomato-expressing astrocytes. **(A)** Low-magnification (2.5x) overview of the entire grid acquired with an external cryo-confocal microscope, showing an overlay of brightfield and tdTomato fluorescence channels with a (**B**) 100x validation of an astrocyte-specific signal. The dashed green rectangle indicates the region of interest (ROI). **(C)** SEM overview of the same grid with an overlay of tdTomato signal acquired with iFLM. The solid green rectangle highlights the ROI corresponding to (**A**). (**D**) View of the ROI after front surface opening overlaid with an iFLM signal from tdTomato validating an ROI. (**E-G**) Higher magnification images used for ROI localization: (**E**) cryo-FLM, (**F**) SEM (pFIB), and (**G**) merged overlay confirming precise targeting. **(H)** Segmented NVU compartments within the acquired cryo-FIB/SEM volume and morphologically correlating with lumen (blue), endothelial cells (yellow), and pericytes (purple) of a cortical blood vessel. (**I, J**) Quantitative comparison of features (lumen diameter, endothelial thickness, pericyte length and lumen area) obtained by cryo-FIB/SEM tomography against previously reported r.t. EM data. Scale bars are (**A, C**) 500 µm, (**B**) 5 µm, (**D**) 100 µm, (**E-H**) 10 µm.

### ROIs definition for subsequent cryo-FIB/SEM volume-EM imaging of an NVU

To validate the physiological relevance of our outputs placed in context, we performed targeted cryo-FIB/SEM volume-EM imaging **(Figure 3)**. As a model target, we have chosen the NVU because it contains several biological elements with known morphology. To precisely locate the NVU compartments within the thick specimen, we utilized the KI mouse model with astrocytes expressing tdTomato fluorescent protein. We extracted the mouse cortex, validated the presence of corresponding ROIs within the extracted region via r.t. light microscopy, and placed the corresponding biopsy punch on the cryo-EM grid (**Supplementary Figure 1**). After soaking the biopsy in 10% PVP with 0.3 M trehalose and subsequent PF, we screened our samples with an external cryo-FLM (**Figure 3A**) and identified 16 unique ROIs within one tissue slice that represent a characteristic tube-like fluorescent signal, consistent with astrocyte endfeet wrapping around vasculature **(Figure 3B**). ROIs were validated with an integrated fluorescence module (iFLM) before (**Figure 3B**) and after (**Figure 3C, D**) the opening of the sample surface. Of note, the NVU is easily identifiable in SEM imaging as a clearly visible lumen surrounded by high-contrast membranes of endothelial cells (**Figure 3E**) and a typical pericyte. We have collected data from a volume of about 32,000 um^3^ that includes a pericyte, endothelial layer, three astrocytic endfeet, and a vascular lumen (**Figure 3G, Supplementary videos 7-8**).

### Physiological relevance of the cryo-FIB/SEM volume-EM imaging outputs

We have further assessed the effects of cryoprotectants on the physiological relevance of our outputs by comparing the morphological features of known NVU subcompartments (**Figure 3H, I**). While the overall architecture of an NVU resembles previously reported ones and contains all the major cellular entities involved in the NVU formation and function (Iadecola, 2017), we have detected significant expansion of the corresponding cellular components. Our results are in line with the previously reported expansion of the biological entities vitrified in the solution with 2-fold higher than normal aCSF osmolarity in comparison to r.t. EM data (Glynn et al., 2024). Specifically, we observe an ∼3-fold and ∼1.4-fold expansion of endothelial layer thickness and pericyte length respectively in comparison to previously reported values (Bonney et al., 2022; Hayden et al., 2018; Lindahl et al., 1997; Nahirney and Tremblay, 2021; Wolburg et al., 2009). For lipid-rich astrocyte endfeet, we see a ∼4-fold expansion that correlates with significant alterations of a largely membranous endothelial layer within NVU. We explain such a drastic difference by the near-native aqueous environment in our case versus the substitution of the water with chemical fixatives often used in r.t. EM which are particularly damaging for extracellular and lipid-rich regions (Korogod et al., 2015).

In the case of the vascular compartment, observed expansion and diversity of vascular units complicates unbiased feature identification. Definition of any unit of vasculature is dependent on proximity to the aorta or vena cava and branching from a higher-order unit. The average diameter of a murine vascular unit is approximately 4 microns for a capillary (Stefanovic et al., 2008; Steinman et al., 2017) and despite the relatively large collected volume of ∼32,000 um^3^, even larger volumes are required to image those vascular units fully. In line with r.t. EM estimations, and when accounting on an observed expansion, the imaged vasculature correlates with a capillary (**Fig. 3H, I**). We also detect a branch in the vascular tree (**movie S7**) that we assign as a capillary cross section.

In summary, the size deviation seen in vitrified samples is either similar or expanded in comparison to r.t. EM data, suggesting that our sample vitrified with cryoprotectants is not altered or squeezed but rather preserved in its physiologically relevant state.

### ROIs definition for subsequent cryoET imaging of astrocyte processes

To validate the physiological relevance of our outputs on subcellular level we performed cryoET imaging of the astrocyte processes (**Figure 4**). Astrocyte processes are a relatively rare feature within general 7,770-fold lower density than synapses (Tsai et al., 2009; Santuy et al., 2020) thus making them a non-trivial target for cryoET. While bead-based cryo-FLM targeting for cellular samples is well-established(Arnold et al., 2016), alternative target-dependent strategies must be utilized for thicker specimens. Unspecific small molecule fluorescent labelling typically generates sufficient signal to navigate with iFLMs, which are typically widefield and produce low-to moderate-quality outputs. So far there are no reliable iFLM data for the translationally relevant KI animal models. For our samples, we employed an external cryo-confocal setup equipped with a superresolution detector (**Figure 4A, B**). Similarly to the cryo-FIB/SEM volume-EM targeting (**Figure 3A, B**), we utilized coordinates of corresponding ROIs (**Fig. 4A, B**) within a biopsy vitrified with a cryoprotectants mixture. After targeted lamellae generation (**Figure 4C, D**), cryoET data were acquired within the regions containing characteristic morphological features of processes (Steinman et al., 2017) followed by a post-TEM validation of the fluorescent signal presence within the imaged lamellae (**Figure 4E-G**). While cryoET data acquisition is expected to alter the fluorescent signal, corresponding tomograms collected from the fluorescent regions correlate with the astrocyte process-specific subcellular morphology (**Figure 4H, Figure 5, Supplementary video 9**).

**Figure 4.**
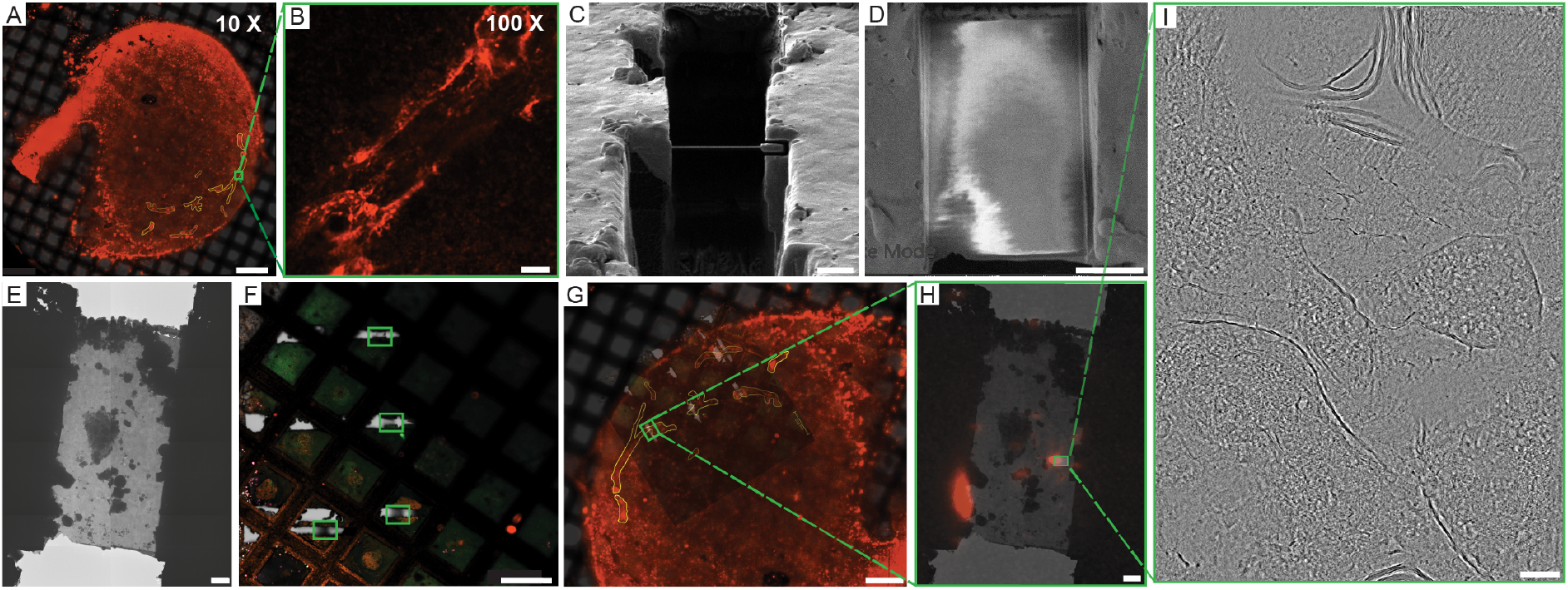
Cryo-FLM guided targeting of astrocytes and lamella preparation via FIB milling. (**A**) Cryo-confocal map of a plunge-frozen mouse cortical tissue with tdTomato-expressing astrocytes (red). Astrocytes are contoured in yellow with a representative fluorescence trace shown on (**B**). (**C**) ROIs-containing lamellae in SEM (**C**), IB (**D**), and cryo-TEM (**E**). Post-TEM cryo-confocal lamellae (**F**, brightfield) realigned to an initial map placing lamellae within the target ROIs (**G**). Post-TEM tdTomato signal (**H**) for the central slice of the collected tomogram (**I**). Scale bars for (**A**) 200µm, (**B-D**) 5µm, (**E, H**) 2 µm, (**F, G**) 100 µm, (**I**) 100 nm.

**Figure 5.**
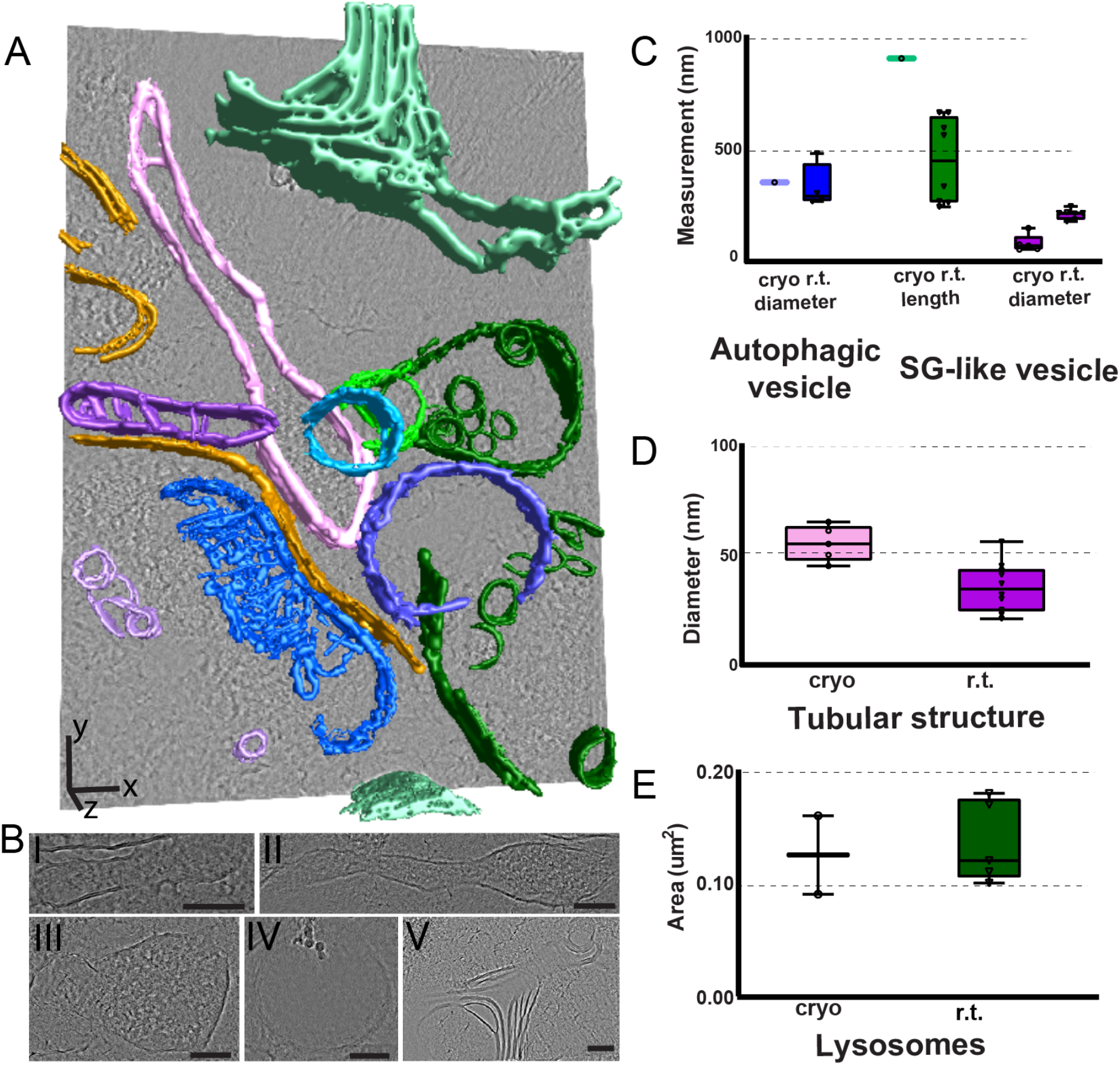
Segmented cryoET imaging data and comparison of subcellular organelles within astrocyte processes. (**A**) Segmentation of a representative tomogram showing distinct subcellular structures: tubular structure (purple), secretory granule (SG)-like vesicle (pink), lysosome (dark green), autophagic vesicle (majorelle blue), mitochondria (blue), peroxisome (light green), vesicle (light purple), unclassified double membrane structures (bronze) and myelin debris (mint). (**B**) The individual structures indicated in (**A**): (I) tubular structure, (II) secretory granule (SG)-like vesicle, (III) lysosome, (IV) autophagic vesicle, and (V) myelin debris. (**C–E**) Quantitative comparison of the dimensions of selected organelles within tomogram and previously reported r. t. EM studies. Scale bars: (**A**) x,y −100 nm, z – 200 nm.

### Physiological relevance of cryoET outputs

Currently, cryoET is capable of obtaining data with a higher resolution than cryo-FIB/SEM volume-EM imaging, thus allowing for visualization of subcellular organelles as well as protein machinery in more detail. We have collected the high-resolution cryoET data from a region exhibiting fluorescent signals of astrocytes’ processes. Among the numerous apparent features, we observed the elongated mitochondria (**Figure 5A**, blue) locked inside an autolysosome-like single layer membranous compartment (**Figure 5A**, bronze) and secretory granules (SG)-like vesicular structures (**Figure 5A**, pink). While mitochondria are ubiquitous and not exclusive for astrocytes in the mouse cortex, elongated SG-like vesicles are found in astrocytes, albeit much less understood (Hur et al., 2010). SGs-like vesicles are densely packed with signaling molecules and they appear as feature-dense elongated subcellular structures in our data. In comparison to previously reported SG structures obtained at r.t. EM, our outputs are much longer and have a 2.6-fold narrower diameter. The localization of the reported SG-like vesicles is unassigned, however the observed morphological differences can be attributed to processes-specific imaging in our case. Elongated SG-like vesicles’ morphology also correlates with its proposed function to navigate the narrow astrocyte processes while keeping them mobile. We also detected peroxisome, autophagic vesicles, and lysosomes for which the size correlation is ambiguous due to the variety of the reported r.t. EM features and lack of the astrocyte processes-specific data. Additionally, the size of these features is heavily dependent on their stage and reported morphology varies significantly. However all our measurements (**Figure 5C-E, Supplementary Figure 7**) were within the reported range, thus validating physiological relevance of the cryoET outputs. At the same time, similar to our cryo-FIB/SEM volume-EM data (**Figure 3**) and previous reports for synaptic vesicles (Glynn et al., 2024), we observed a minor 1.6-fold expansion for the tubular astrocyte-specific structures (**Figure 5A**, purple). Interestingly, we also detected the lipid-rich membranes that we identified as myelin debris (**Figure 5A**, green) due to the high contrast, its irregular shape, and intracellular location. The proximity of lysosomes and autophagic vesicles confirms this hypothesis and defines the imaged processes as actively degrading the internalized myelin sheath. Additionally, proximity of peroxisomes correlates with active lipid degradation and metabolic support. Taken together, we further confirm the physiological relevance of our outputs and define cryoET as a comparable but more physiologically relevant imaging modality as opposed to conventional r.t. EM.

## Discussion

Here we benchmarked PF on mouse brain tissues and subsequent lamellae preparation (**Figure 1**) and validated the physiological relevance of the corresponding cryo-imaging outputs. In our hands, tissue slices of up to ∼100 microns can be vitrified robustly however this limit may be different for each organ and corresponding cryoprotectants mixture.

Based on the data described on **Figure 2**, we have defined 10% PVP with 0.3 M trehalose as the most appropriate cryoprotectant mixture. In line with previous reports (Glynn et al., 2024), a 2-fold osmolarity increase with 10% PVP and 0.3 M trehalose does not result in visible artifactual features when the samples are soaked in cryoprotectant-containing solution for up to 30 minutes and therefore can be utilized for the low-resolution morphological assessment. However, due to the potential interference of trehalose with native protein conformation, we suggest using PVP-based mixtures for the high-resolution structural determination. While there are no reports about possible interference of PVP with the protein structures, it does not contain the hydroxyl groups that might substitute the water shell around the proteins as opposed to several hydroxyl groups in trehalose.

In our case, we were targeting regions of 5 microns thick for which we detected the fluorescence signal from astrocyte processes (**Figures 3-5**). In cases where more precise lamellae-level targeting is required, our strategy would need further optimization. For example, precise targeting within thicker translationally relevant specimens would imply crossing several KI lines (Matsui et al., 2024). In addition, further development in superresolution cryo-FLM (Dahlberg et al., 2020; Dahlberg and Moerner, 2021) and live fluorescent monitoring with tri-coincident setups (Boltje et al., 2022; Sica et al., 2024; Perton et al., 2025) will significantly simplify the subcellular targeting with a 100-200 nanometers precision and retain the features of interest within a lamella.

While the exact mechanism behind the successful PF-assisted vitrification is unclear, we suggest that in the case of brain tissue, the presence of natural cryoprotectants is the major driving force behind successful vitrification. With at least the liver and heart typically being more cryoprotectant-rich than the brain, we expect that those organs can also be vitrified after careful optimization of the cryoprotectants solution. On the other hand, the brain is tightly packed with proteins and other features as opposed to alternative samples where features are located sparsely (Pöge et al., 2025) making such samples not amenable to successful PF at this point.

Our work successfully benchmarks PF for thicker mammalian brain tissue specimens to obtain artifact-free and physiologically relevant cryo-imaging data. While further studies are required to reveal the details of why PF is capable of freezing such thick specimens, we speculate that extremely protein-dense and endogenous cryoprotectant-rich brain tissue are the perfect model samples for PF. This assumption suggests that after meticulous optimization of the chemical cryoprotectant mixture, other organs like liver and heart are potentially subjectable to efficient PF as well. We have also detected a previously noted expansion of the biological features even in the cryoprotectant mixture with the 2-fold higher osmolarity than typical aCSF. Most importantly, our pipeline establishes a solid basis for future cryo-imaging studies using basic vitrification instrumentation that is generally available in laboratories applying cryogenic electron microscopy at multiple scales.

## Acknowledgments

We thank Zephan Melville and Edward Eng for the help with the data collection as per NCCAT proposals ID BAGOK250359 and BAGOK221001. We also thank Richard G. Held and Tamara Basta for fruitful scientific discussions about vitrification, Jason Pugh for providing access to an osmometer, Naomi Sayre for the kind donation of the CX30-CreERt2 mice, and UTHSCSA Mouse Genome Engineering and Transgenic Facility for the help with animal experiments as well as UTHSCSA cryo-EM facility for providing an access to the plunge freezing setup.

OK is supported by The Robert A. Welch Foundation research grant #AQ-2197-20240404 and The Voelcker Foundation Young Investigator Award #10012268. Some of this work was performed at the National Center for CryoEM Access and Training (NCCAT) and the Simons Electron Microscopy Center located at the New York Structural Biology Center, supported by National Institutes of Health (Common Fund U24 GM129539, NIGMS R24 GM154192), the Simons Foundation (SF349247) and NY State Assembly. A portion of this research was supported by National Institutes of Health grant U24 GM139174 and performed at the CU Boulder Center for Cryo-ET (CCET) at the University of Colorado, Boulder. A portion of this research was supported by National Institutes of Health grant U24 GM139166 and performed at the Stanford-SLAC CryoET Specimen Preparation Center (SCSC) at Stanford-SLAC.

## Notes

### Competing Interest Statement

The authors have declared no competing interest.

