## Supplementary Figures and Tables for "Cryoprotectants-assisted plunge freezing of thick brain tissue specimens for targeted physiologically relevant cryo-imaging *in situ*"

**Supplementary Figure 1**. Schematic representation of the pipeline for cryoET imaging of mouse brain tissue specimens. After extraction of the tissue of interest (here - mouse brain), fresh tissue is sliced with a vibratome setting of 30-100 microns. If present, ROIs are confirmed at room temperature and 1.5 mm biopsy punches containing ROIs are prepared. Next, tissue slices are positioned centrally on the cryo-EM grids which are placed in a transfer container (here, an 1.5 mL eppendorf tube) while each sample is soaked in a solution containing cryoprotectants (for 20-60 minutes). Samples are vitrified with a plunge freezing setup that allows one-side blotting (here, Leica EM GP2). Plunge-frozen tissues are screened with an external cryo-FLM setup to confirm the presence of ROIs presence (here, a Zeiss LSM900 with an Airyscan 2 and Linkam cryo-stage). For each ROI, a set of coordinates is obtained for subsequent targeted thick tissue thinning. Thinning is performed in semi-automated fashion with LMIS-cryoFIB/SEM (here, Aquilos 2) or cryo-pFIB/SEM (here, Hydra Bio) for subsequent cryoET data collection.

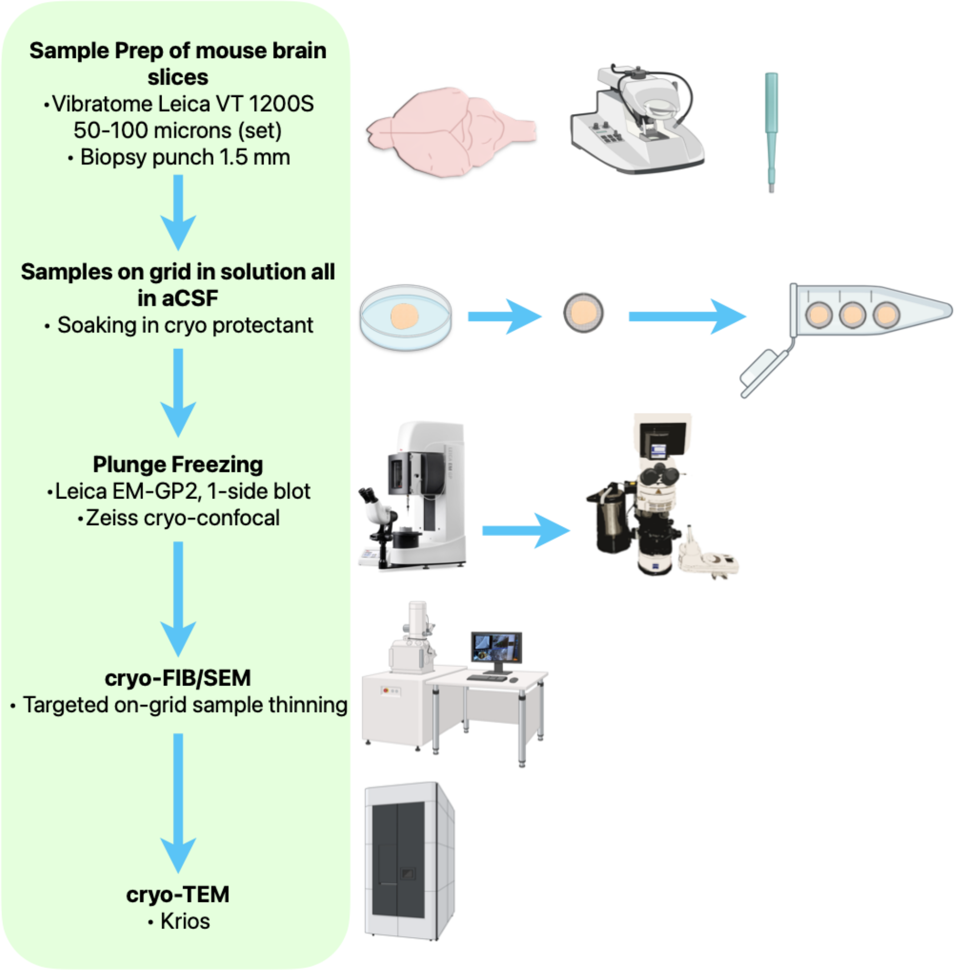

**Supplementary Figure 2.** Representative FFTs obtained with Warp and derived from zero tilts for the tomograms collected on the tissues prepared with cryoprotectants mixtures tested for **Fig. 2**. Experimental contrast transfer function is shown in blue and estimated function is in pink, defocus, and resolution.

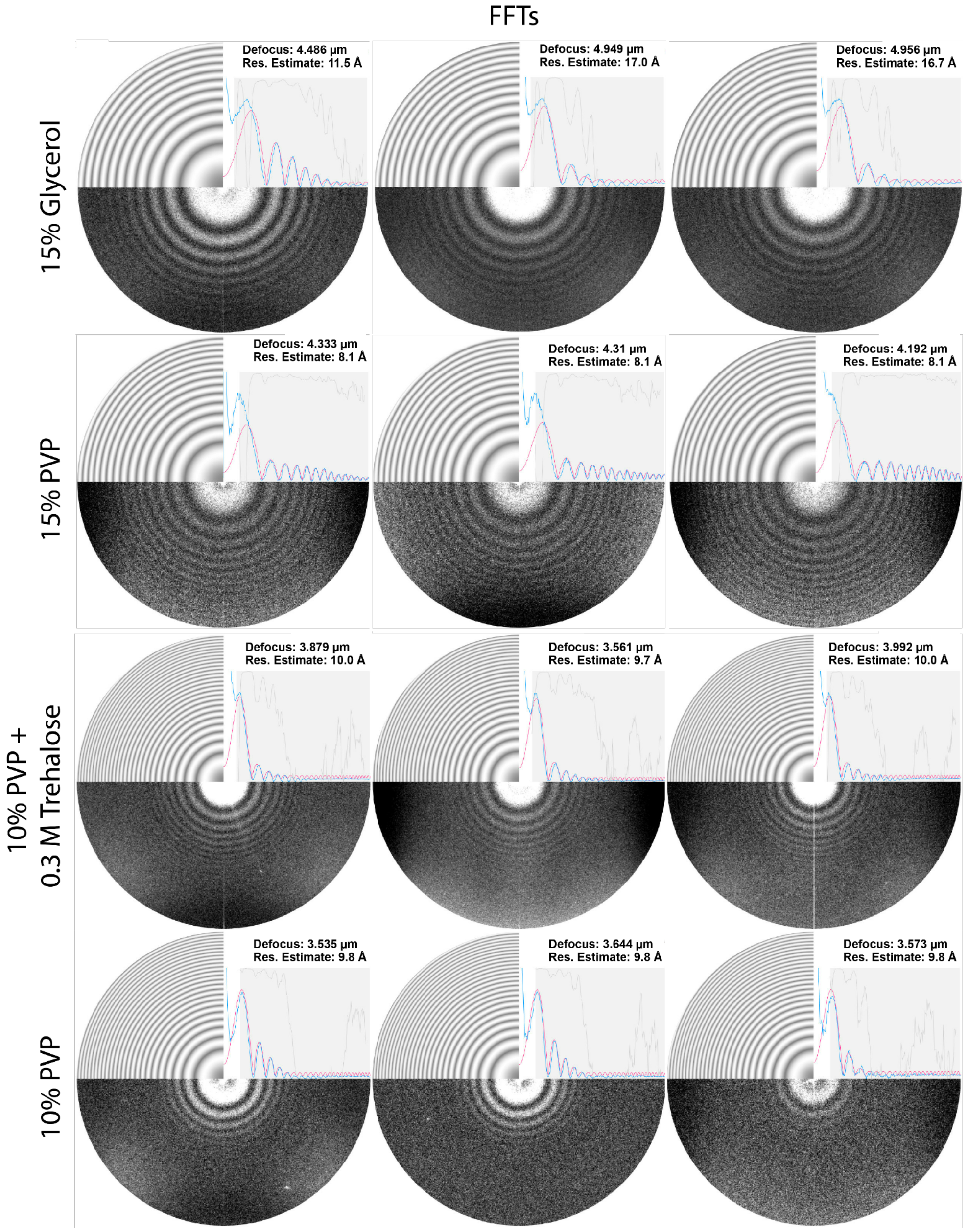

**Supplementary Figure 3.** Representative search maps acquired in cryo-TEM for the samples plunge frozen in 15% glycerol with 4-fold increased osmolarity. Depicted myelin sheaths exhibit artificial morphology that is not reported in the literature and therefore defined as artefactual. Scale bars - 5 μm.

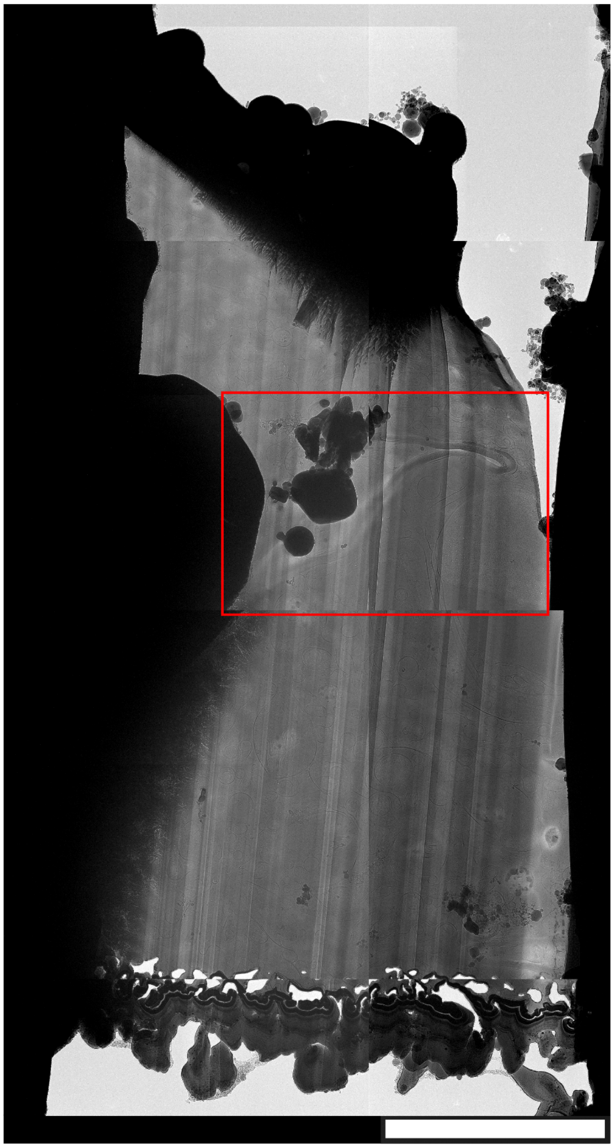

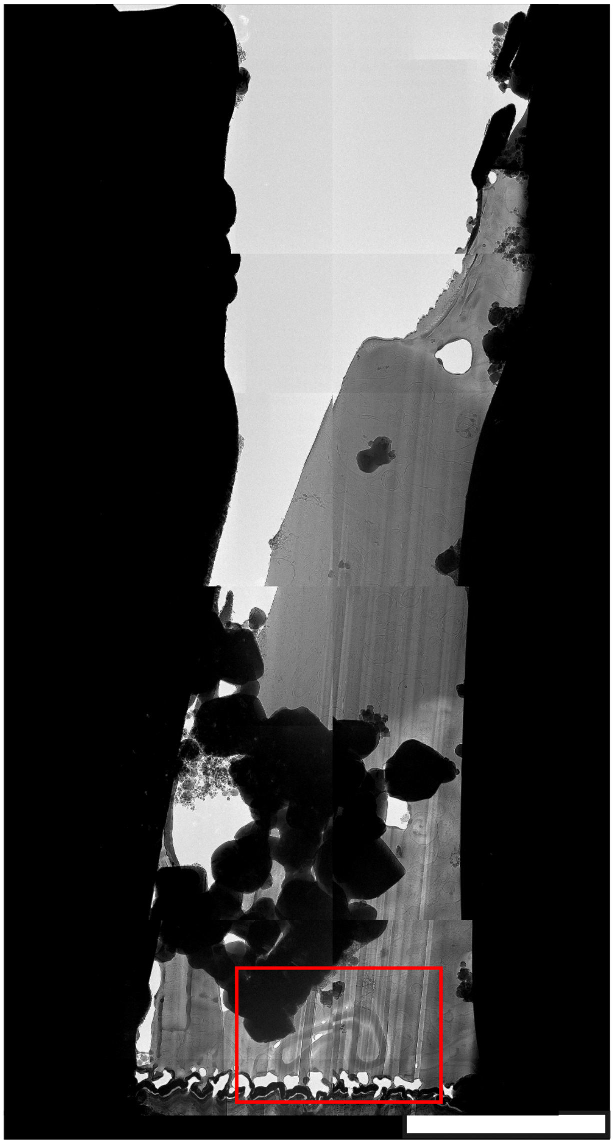

**Supplementary Figure 4**. Representative SEM images during the course of automated thinning with AutoTEM on LMIS-cryoFIB/SEM. Times (minutes:seconds, after the start of each step) and thinning steps are displayed at the top of each image. From left to right, images depict the thinning process from the initial slab of 25-50 μm (top and middle rows) or less (bottom row) to the final lamellae, and are taken after the completion of each thinning step. With LMIS, notches are fusing within the first minutes of Rough milling and thus should be refreshed before or after polishing to maintain the stress relief.

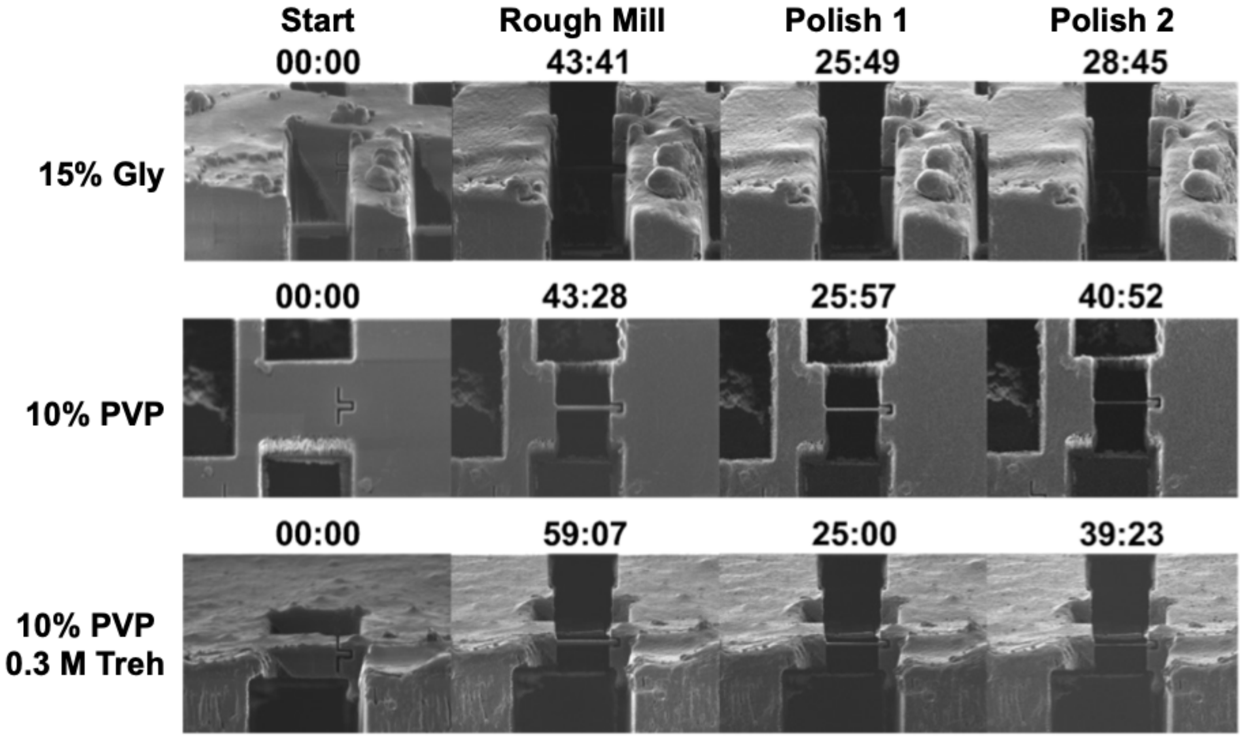

**Supplementary Figure 5.** Cryo-FIB/SEM volume imaging outputs processing with DragonFly. After initial alignment and images filtering, curtaining is removed from aligned stacks. (**A**) Argon milling resulted in a heavily charged set of slices which were split from the entire volume collected and required a set of dedicated algorithms to retrieve the meaningful information. In contrast, applying Xenon (**B**) resulted in much less severe charging artifacts and required much less postprocessing.

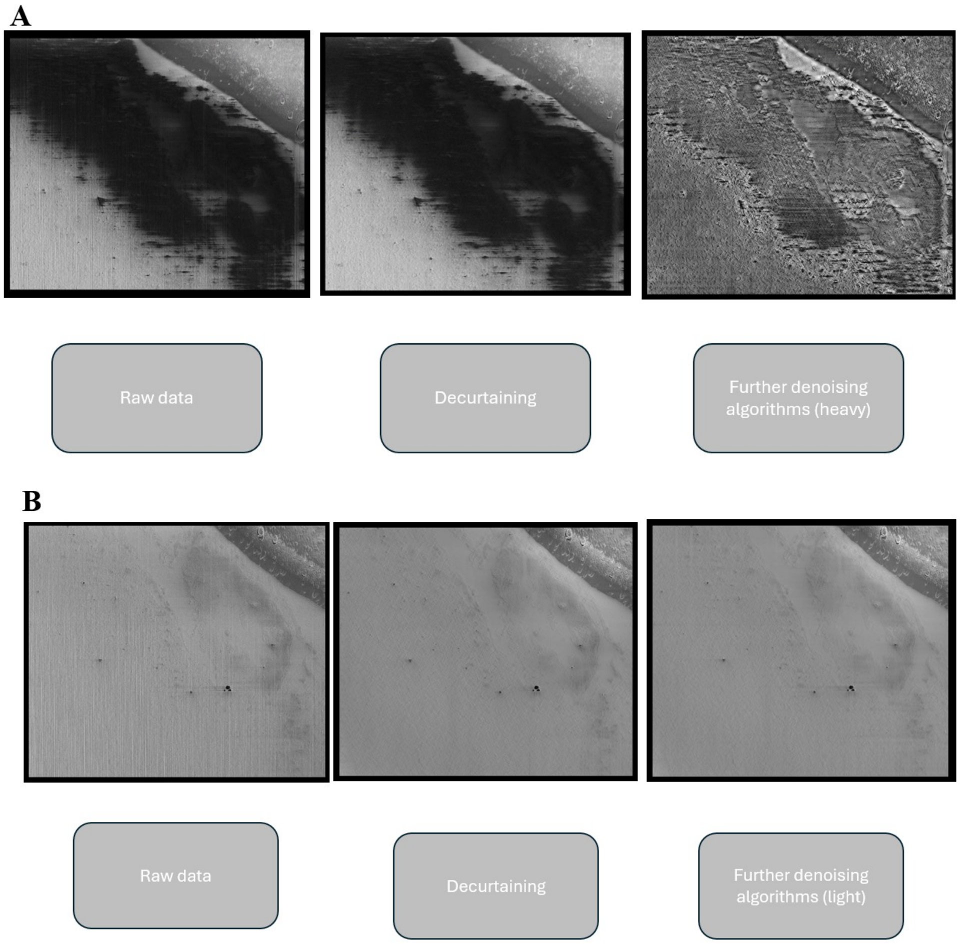

**Supplementary Figure 6**. Additional features identified within the tomogram on Figure 5A as described in the Materials & Methods section. (**A**) Heavily distorted mitochondria located within the putative lysosomal compartment and is potentially undergoing degradation. (**B**) Peroxisome bound to unidentified membranous feature. Scale bars - 100 nm.

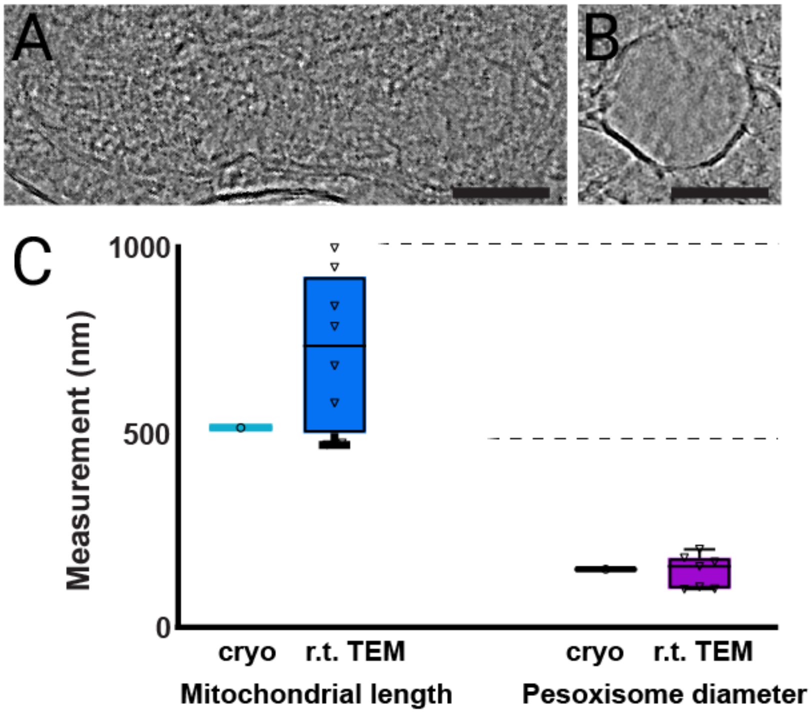

**Supplementary Table 1** with cryo-confocal acquisition settings for Zeiss LSM 900.

| **Parameters** | 10 x | 100 x |
|---|---|---|
| **Scan Mode** | Frame | Frame |
| **Scan Zoom** | 1.0 | 1.3 |
| **Rotation** | 0° | 0° |
| **Sampling** | 0.64 | 3.31 |
| **Pixel Time** | 0.76 µs | 2.06 µs |
| **Frame Time** | 1.86 s | 10.13 s |
| **LSM Scan Speed** | 8 | 6 |
| **Scan Direction** | Bidirectional | Bidirectional |
| **Line Step** | 1 | 1 |
| **Averaging** | 2 | 4 |
| **Averaging Mode** | Line | Line |
| **Averaging Method** | Mean | Mean |

**Supplementary Table 2** with the AutoTEM Cryo settings for automated lamellae generation.

| **Auto-TEM step** | **Current (pA)** | **Offset (um)** | **Pattern Overlap (%)** | **Depth correction (%)** |
|---|---|---|---|---|
| Rough | 7000 | 12.5 | N/A | 50 |
| Medium | 3000 | 8 | 500 | 150 |
| Fine | 1000 | 0.8 | 300 | 100 |
| Finer | 300 | 0.4 | 300 | 100 |
| Pol1 | 30 | 0.15 | 300 | 90 |
| Pol2 | 10 | 0 | 300 | 90 |

**Supplementary Videos.**

**Supplementary Video 1.** Full tomogram for Figure 1K. Scale bar - 100 nm.

**Supplementary Video 2.** Full tomogram for Figure 1K. Scale bar - 100 nm.

**Supplementary Video 3.** Full tomogram for Figure 2B (15% glycerol). Scale bar - 100 nm.

**Supplementary Video 4.** Full tomogram for Figure 2C (10% PVP + 0.3 M trehalose). Scale bar - 100 nm.

**Supplementary Video 5.** Full tomogram for Figure 2D (10% PVP). Scale bar - 100 nm.

**Supplementary Video 6.** Full tomogram for Figure 2E (15% PVP). Scale bar - 100 nm.

**Supplementary Video 7.** Segmented NVU within the acquired cryo-FIB/SEM volumes with Xe.

**Supplementary Video 8.** Segmented NVU within the acquired cryo-FIB/SEM volumes with Ar.

**Supplementary Video 9.** Full tomogram for Figure 4I (10% PVP with 0.3 M trehalose). Scale bar = 100 nm.
